## Supplementary Figures for "Learning tissue representation by identification of persistent local patterns in spatial omics data"


S1 Kasumi can take as input a vector of marker abundances . . . . . . . . . . . 2
S2 Additional results for the responder group of samples from the CODEX CTCL dataset . . . . . . . . . . . . . . . . . . . . . . . . . . . . . . . . . . . . . . . 3
S3 Additional results for the non-responder group of samples from the CODEX CTCL dataset . . . . . . . . . . . . . . . . . . . . . . . . . . . . . . . . . . . 4
S4 Predictor-target importances of CTCL relevant Kasumi clusters . . . . . . . . 5
S5 Additional results for the sensitive group of samples from the IMC BC dataset 6
S6 Additional results for the resistant group of samples from the IMC BC dataset . 7
S7 Distribution of the Adjusted Rand Index (ARI) across samples . . . . . . . . . 8
S8 Quantification of the value added by the inclusion of the paraview accounting for marker interactions in the IMC BC dataset and predictor-target importances 9
S9 Parameter sensitivity analysis . . . . . . . . . . . . . . . . . . . . . . . . . . 10
S10 Computational complexity of Kasumi . . . . . . . . . . . . . . . . . . . . . . . 10
S11 Application of Kasumi on PDAC SMI (pre-commercial CosMx) transcriptomics data . . . . . . . . . . . . . . . . . . . . . . . . . . . . . . . . . . . . . . . . . 11

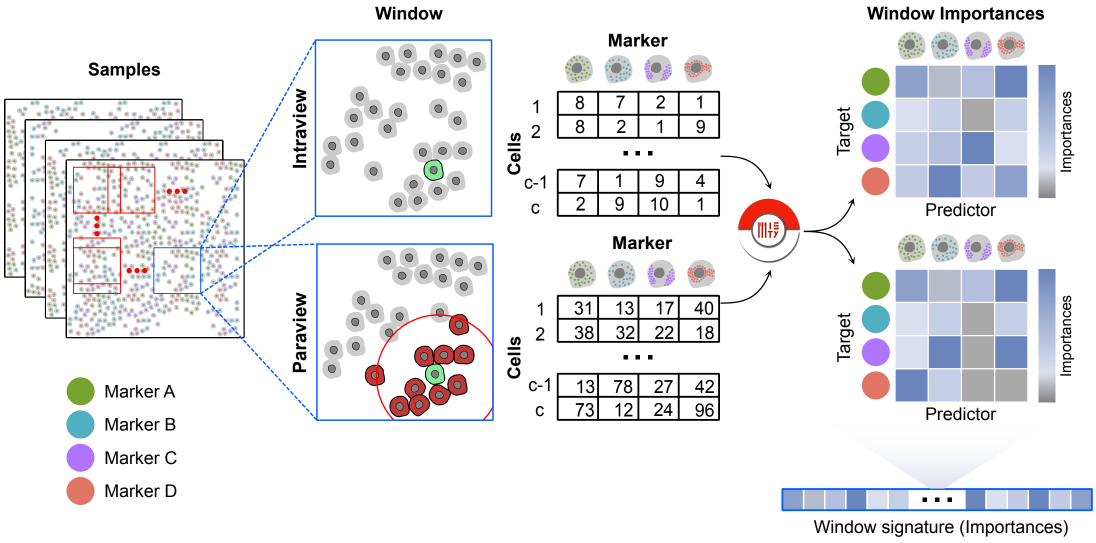


**Supplementary Figure S1:** Kasumi can take as input a vector of marker abundances. Unlike the scenario in which input is provided at the level of cell types (Figure 1), predictor-target relationships are also extracted from the intraview to form a window signature and consequently the first Kasumi representation.

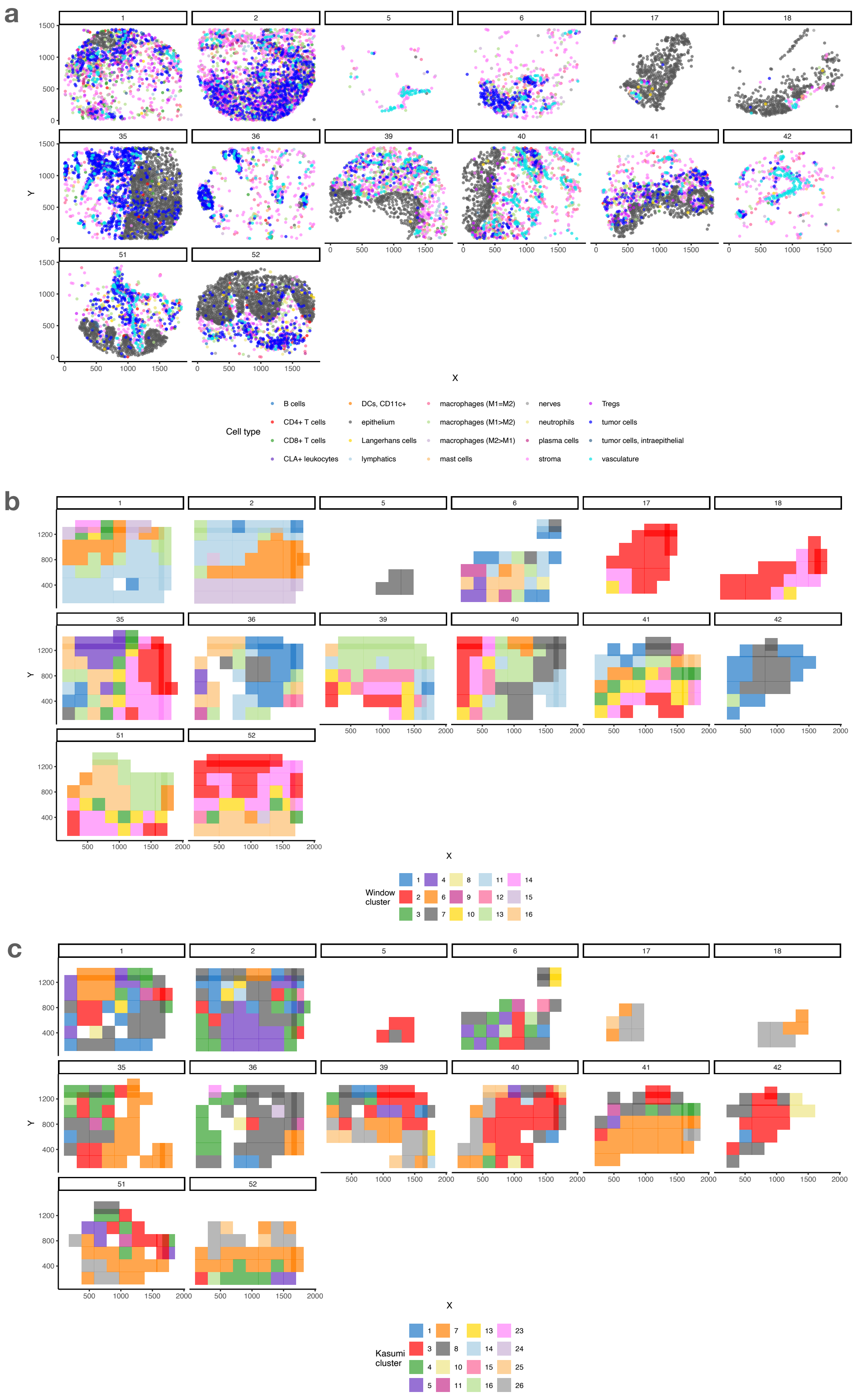


**Supplementary Figure S2:** Additional results for the responder group of samples from the CODEX CTCL dataset. a) Spatial locations of all cell types, b) the cell-type-composition-based window clusters (WCC) and c) cell-type-relationship-based Kasumi clusters.

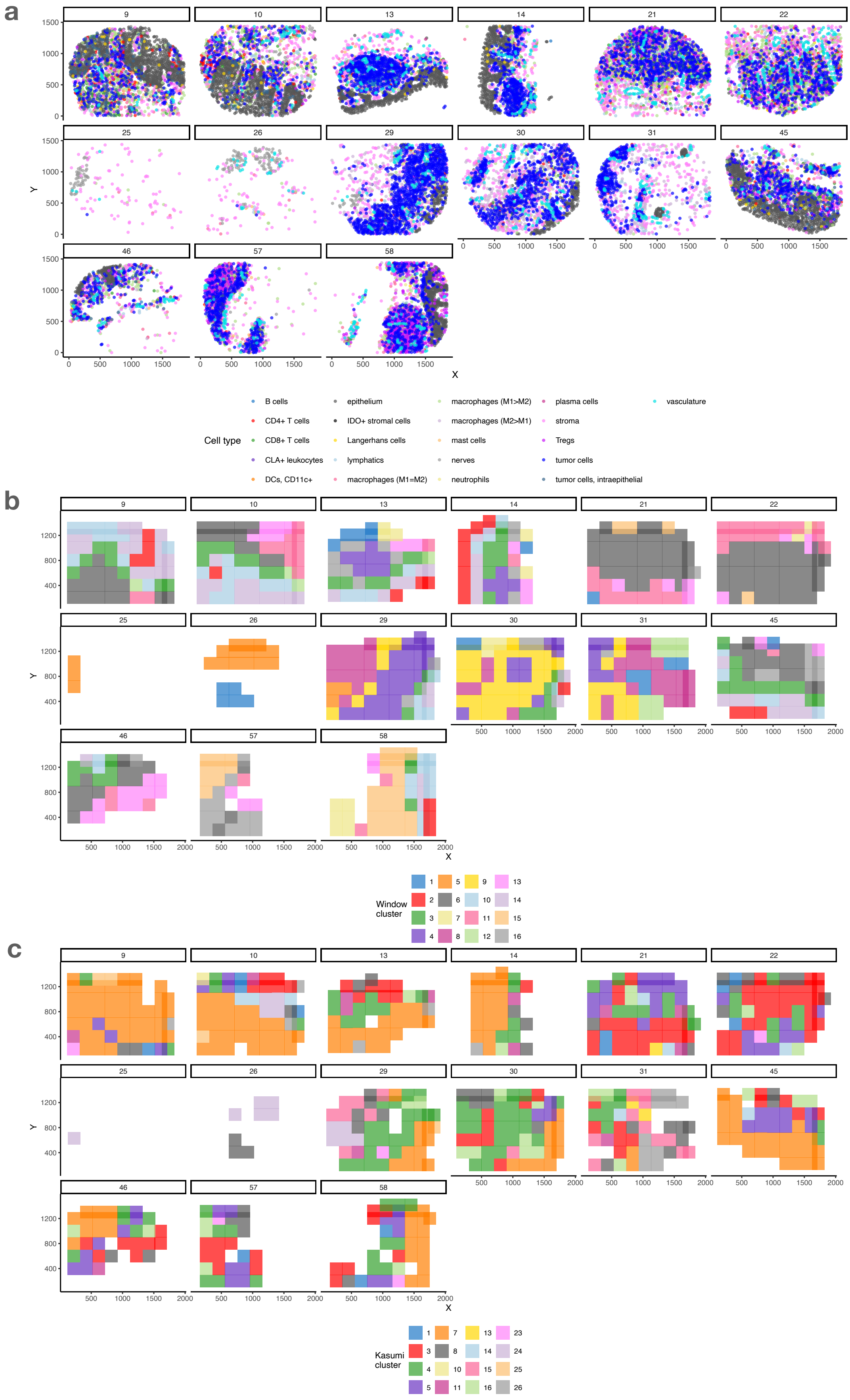


**Supplementary Figure S3:** Additional results for the non-responder group of samples from the CODEX CTCL dataset. a) Spatial locations of all cell types , b) the cell-type-composition-based window clusters (WCC) and c) cell-type-relationship-based Kasumi clusters.

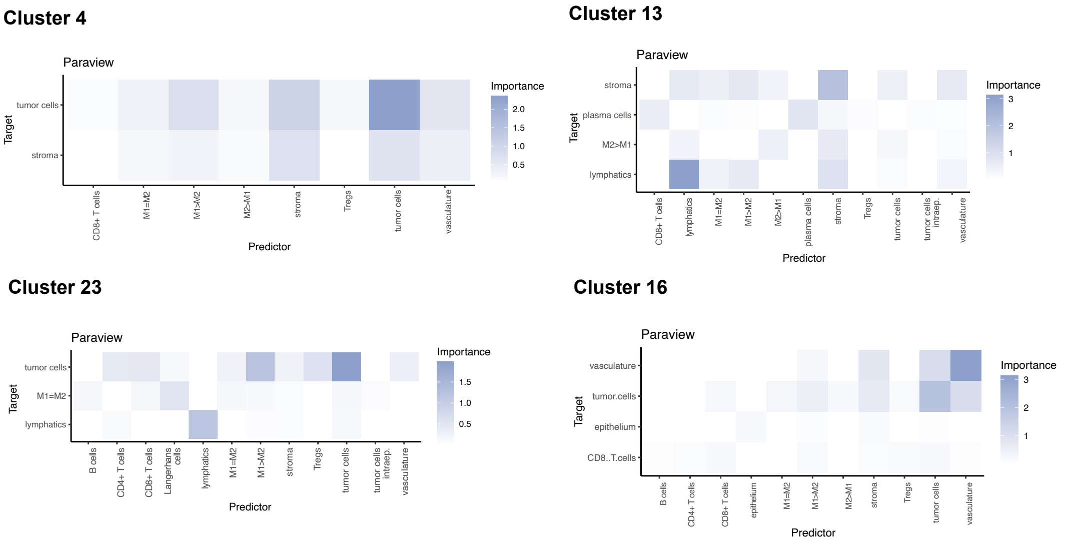


**Supplementary Figure S4:** Predictor-target importances of CTCL relevant Kasumi clusters. Complete set of feature importances for the most relevant Kasumi clusters in the task of distinguishing responders from non-responders in the cutaneous T cell lymphoma (CTCL) data. Shown are targets for which adding the paraview spatial context resulted in an increase of explained variance of more than 1. All importances below 0 are shown as white. Source data are provided as a Source Data file.

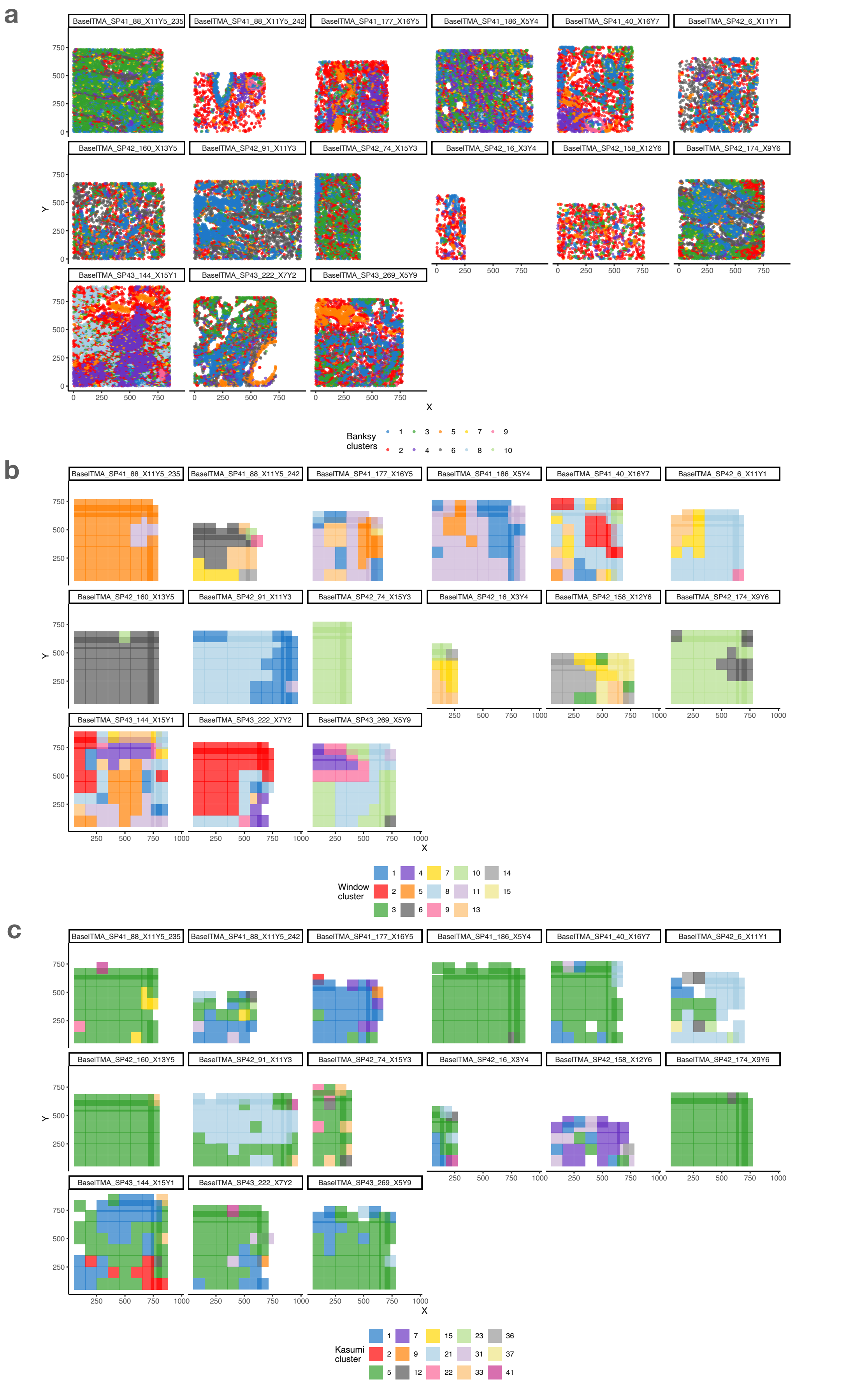


**Supplementary Figure S5:** Additional results for the sensitive group of samples from the IMC BC dataset. a) BANKSY clusters, b) the cell-type-composition-based window clusters (WCC) and c) marker-abundance-relationship-based Kasumi clusters.

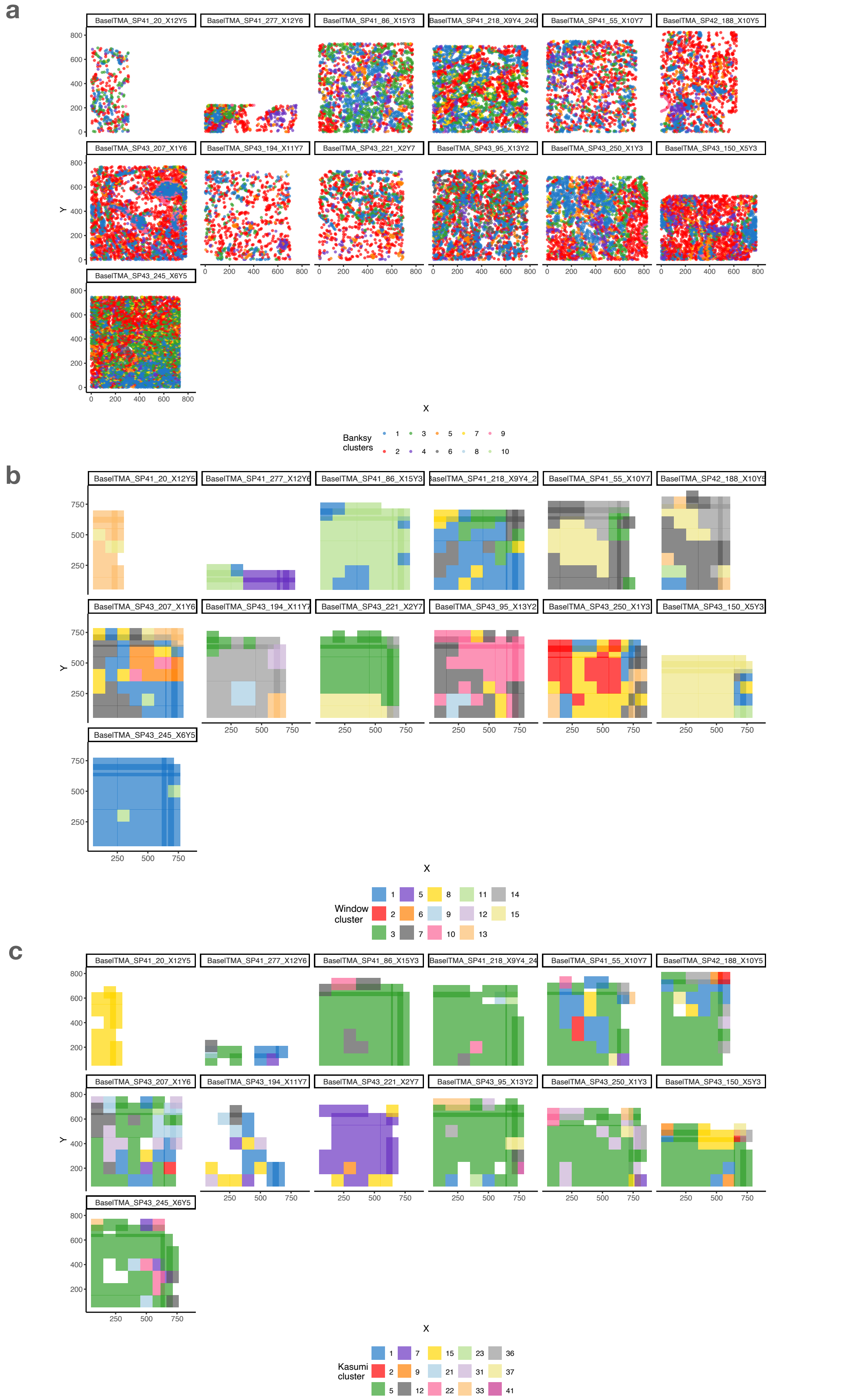


**Supplementary Figure S6:** Additional results for the resistant group of samples from the IMC BC dataset. a) BANKSY clusters , b) the cell-type-composition-based window clusters (WCC) and c) marker-abundance-relationship-based Kasumi clusters.

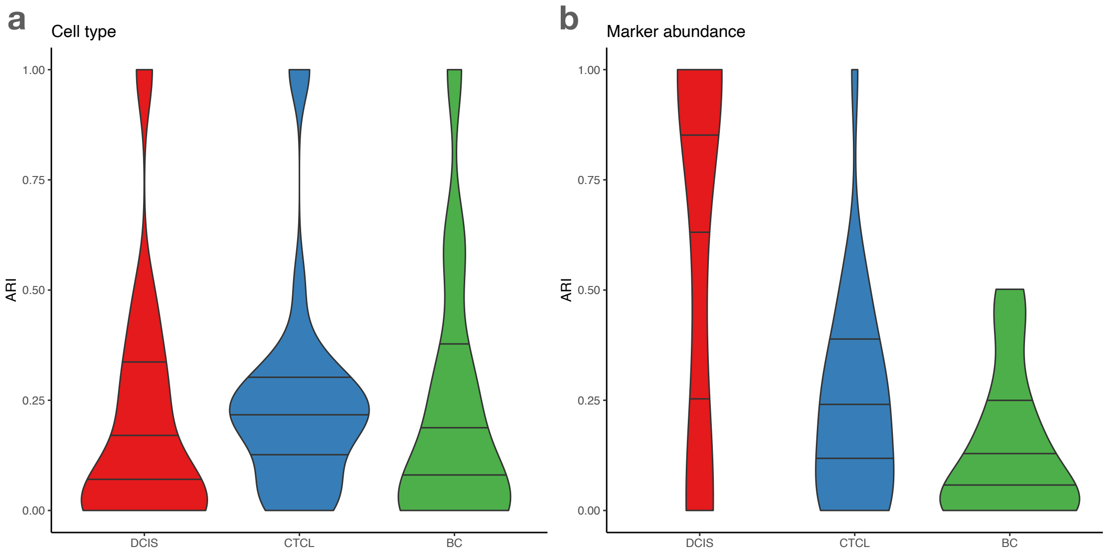


**Supplementary Figure S7:** Distribution of the Adjusted Rand Index (ARI) across samples. Comparison of the Kasumi clustering with WCC on a) cell type and b) marker abundance input per dataset. Horizontal lines mark the 25th, 50th and 75th percentile. Source data are provided as a Source Data file.

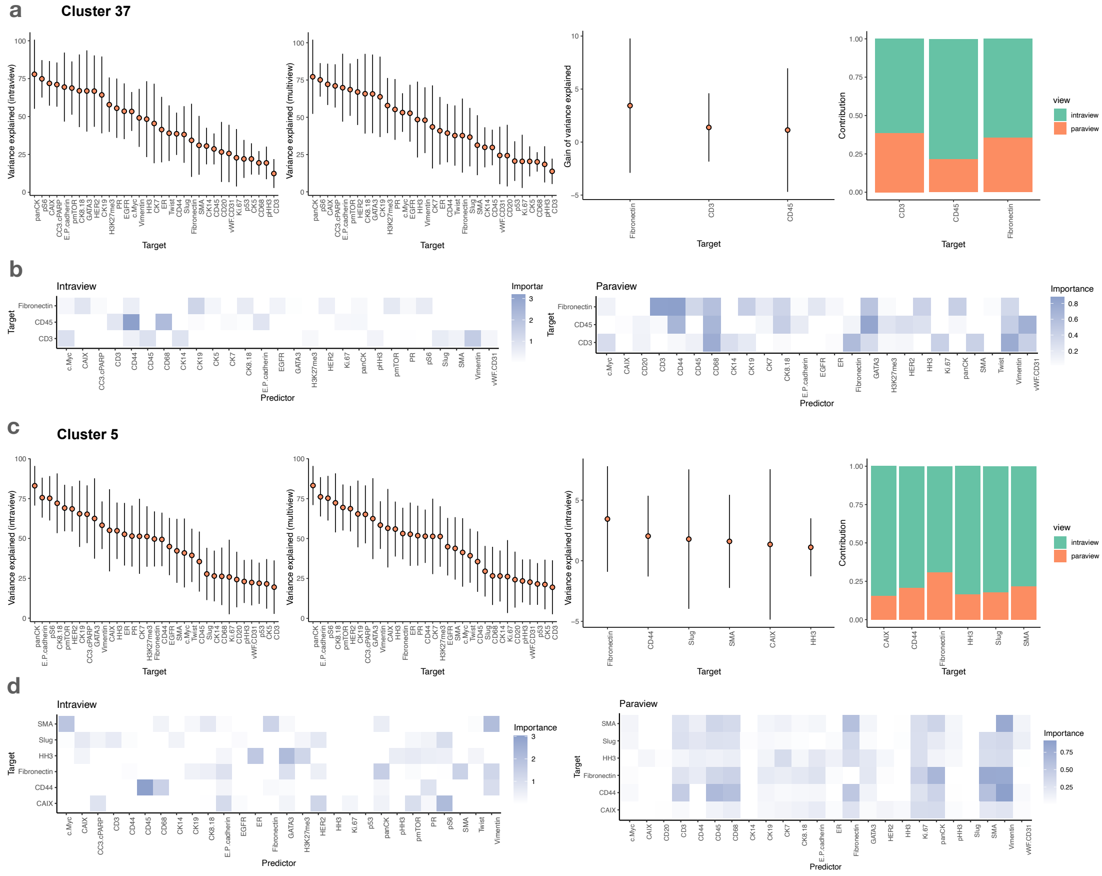


**Supplementary Figure S8:** Quantification of the value added by the inclusion of the paraview accounting for marker interactions in the IMC BC dataset and estimated predictor-target importances. a) Performance and contributions per target for Kasumi marker-abundance cluster 37. Points represent the mean and the bars represent standard deviation across windows labeled with the corresponding cluster (n=7). b) Complete set of intraview and paraview predictor-target importances for cluster 37. c) Performance and contributions for Kasumi marker-abundance cluster 5. Points represent the mean and the bars represent standard deviation across windows labeled with the corresponding cluster (n=804). d) Complete set of intraview and paraview predictor-target importances for cluster 5. Subpanels in a and c display the variance explained per target marker for a model with an intraview only, for a multiview model, the gain of variance explained between the multiview and the intraview model (filtered for targets with gain of more than 1), and relative contribution of the views for the filtered targets. In b and d shown are targets for which adding the paraview spatial context resulted in an increase of explained variance of more than 1. All importances below 0 are shown as white. Source data are provided as a Source Data file.

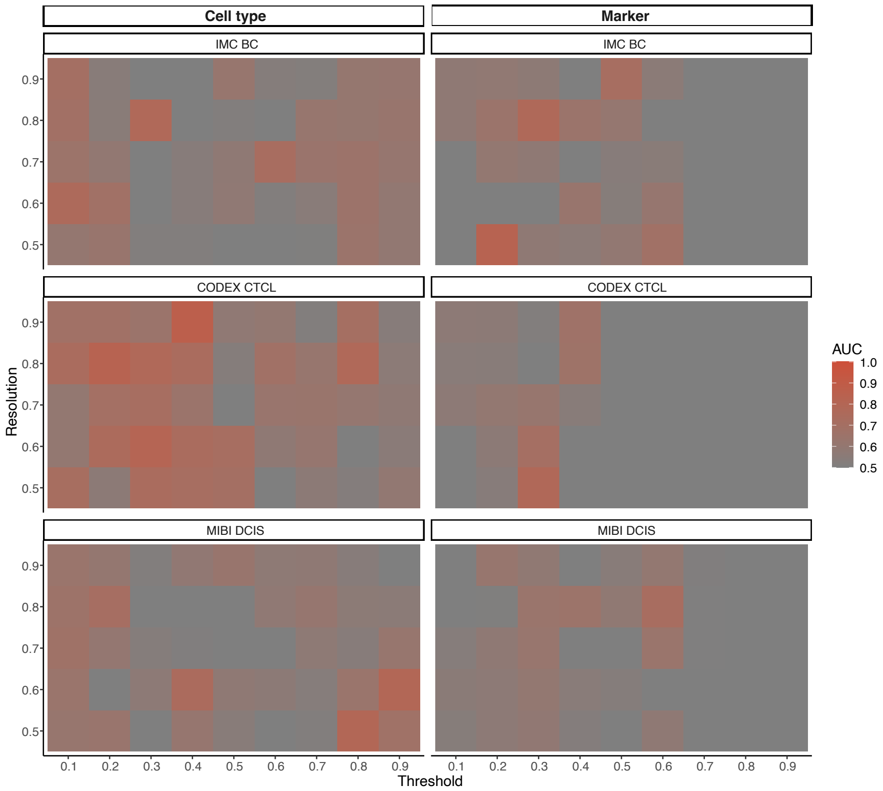


**Supplementary Figure S9:** Parameter sensitivity analysis. Sensitivity of the performance of Kasumi representation to the cutoff threshold for the cosine similarity used to remove edges from the graph of windows and the resolution parameter of the Leiden clustering of that graph. Source data are provided as a Source Data file.

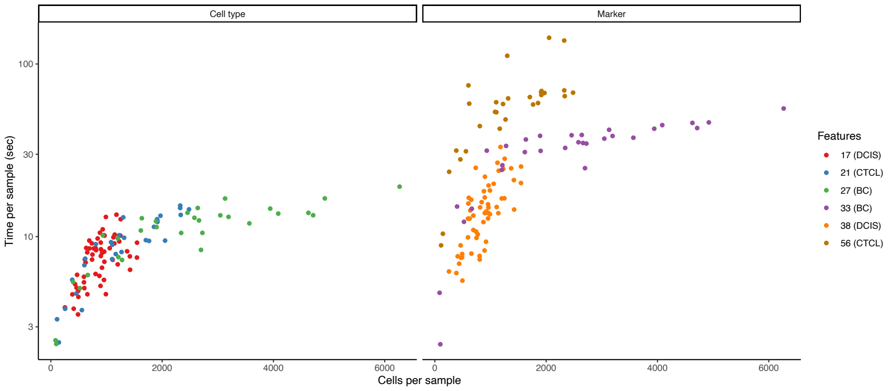


**Supplementary Figure S10:** Computational complexity of Kasumi. Kasumi runtime in seconds per sample as a function of the number of cells and features in that sample. Each point corresponds to a single sample. Source data are provided as a Source Data file.

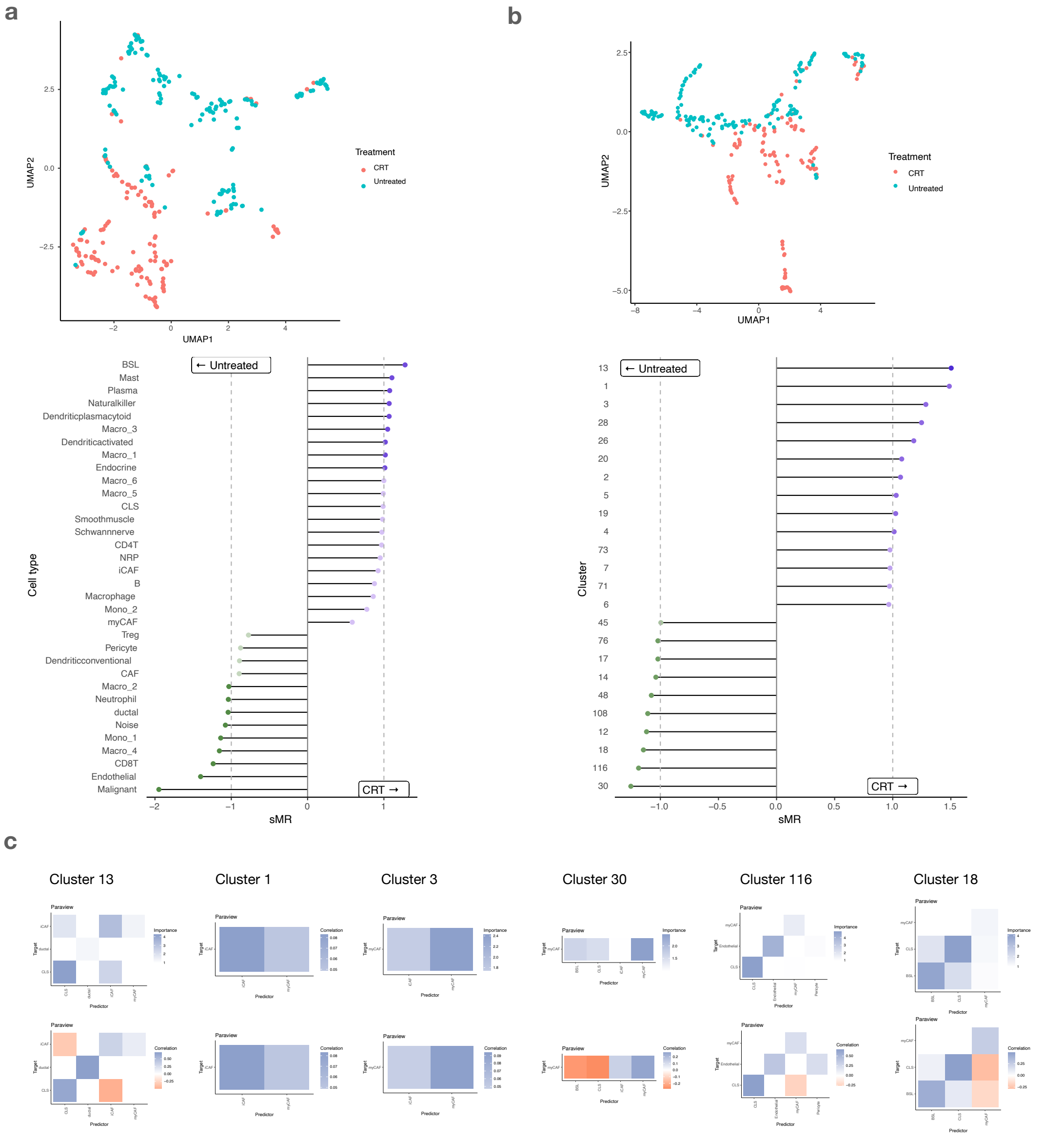


**Supplementary Figure S11:** Application of Kasumi on PDAC SMI (pre-commercial CosMx) transcriptomics data. We selected 300 samples (fields of view) coming from 12 patients. We removed samples from one treated patient that in addition to the capecitabine or 5-fluorouracil (CRT) treatment also received losartan treatment. As input representation we used the most granular cell type information for the more than 700000 cells identified from the expression of a panel of 960 genes. Running Kasumi on all samples on a newer generation laptop with 8 processing cores took in total approximately 85 minutes. a) UMAP and sMR of the representation of the samples by the distribution of cell types. The condition can be predicted without the need for spatial resolution or neighborhood analysis. The non-spatial global composition-based representation already performs very well (AUROC = 0.96). The proportion of basal-like (BSL) malignant cells, Mast, Plasma and other immune cells is highly relevant to the prediction of samples from treated (CRT) patients and the proportion of Malignant and Endothelial cells are relevant to the prediction of untreated samples. b) UMAP and sMR of the Kasumi representation of the samples. Applying Kasumi on the PDAC data resulted in comparably high performance (AUROC = 0.97), but provides a complementary spatial explanation. c) Spatially consistent with the findings presented by Shiau et al., Kasumi identified complex (low-correlation) patterns of cancer associated fibroblasts (inflamatory and myofibroblastic CAF), contiguous ductal structures and absence of inflammatory CAFs in the neighborhood of classical (CLS) malignant cells as most relevant predictors of treated samples. On the other hand, most relevant predictors of untreated samples are Kasumi clsuters are spatially contiguous patterns of BSL and CSL cells lacking involvement of myofibroblastic CAFs. Source data are provided as a Source Data file.
